# Binding mode of isoxazolyl penicillins to a Class-A *β*-lactamase at ambient conditions

**DOI:** 10.1101/2025.05.23.655743

**Authors:** Gargi Gore, Andreas Prester, David von Stetten, Kim Bartels, Eike C. Schulz

## Abstract

The predominant resistance mechanism observed in Gram-negative bacteria involves the production of *β*-lactamases, which catalyse the hydrolysis of *β*-lactam antibiotics, thereby rendering them ineffective. Although isoxazolyl penicillins are available since the 1970s, there are currently no structures in complex with class-A *β*-lactamases available. In order to support the rational development of new *β*-lactamase inhibitors, we have analysed the structure of the clinically relevant *β*-lactamase CTX-M-14 from *Klebsiella pneumoniae* near physiological temperatures. By utilizing serial synchrotron crystallography, we demonstrate the acyl-enzyme intermediates of the catalytically impaired CTX-M-14 mutant E166A in complex with three isoxazolyl penicillins: oxacillin, cloxacillin and dichloxacillin. While the three derivatives differ only by one and two Cl atoms, respectively, they show marked differences in their binding mode.

## 1 Introduction

Bacterial resistance to antibiotics is a leading cause of millions of deaths every year. About 13.6% of the total annual global infection-related deaths are associated with 33 bacteria [1]. Since the discovery of penicillin almost a century ago, antibiotics, especially *β*-lactams, remain irreplaceable. *β*-lactams are the most important as well as the most widely prescribed class of antibiotics against Gram-negative infectious bacteria [2].

Most bacteria have an innate resistance to certain antibiotic groups as a result of low affinity, cell wall impermeability, and absence of antibiotic receptor [3]. Changes to this susceptibility can either be caused by spontaneous chromosomal mutations leading to the acquisition of resistance genes or the development of an extrachromosomal resistance mechanism. As a result of a Horizontal Gene Transfer (HGT), acquired extrachromosomal antibiotic resistance is usually contained within mobile DNA element of the bacteria. Hence, mobile genetic elements like plasmids, conjugative transposons, and integrons are a driving force for bacterial multidrug resistance [4, 5]. Among the numerous bacterial resistance mechanisms, the most prevalent include enzymatic modification of the antibiotic, target site modification, generation of an alternative metabolic pathway, increasing concentration of antagonist of the antibiotic, or active removal of the antibiotic from the cell [3].

The primary mode of resistance in Gram-negative bacteria is enzymatic degradation of antibiotics by *β*-lactamases [2, 6]. The bacterial cell wall comprises heavily cross-linked peptidoglycans. These individual peptidoglycans are produced in the cell, but the cross-linking takes place beyond the cytoplasmic membrane, catalysed by cell-wall transpeptidases. Transpeptidases have an active serine and proceed via an acylation-deacylation pathway to link two peptide strands. *β*-lactam antibiotics effectively inhibit this catalytic cycle due to the stereochemical resemblance of the *β*-lactam with the terminal peptide residues involved in the cross-linking. Thus, the transpeptidases form an irreversible penicilloyl-enzyme complex, thereby blocking peptidoglycan cross-linking during cell wall synthesis, increasing the likelihood of cell lysis and death [2]. Therefore, these transpeptidases are also referred to as penicillin-binding proteins (PBPs) [7].

To avoid this, bacteria produce *β*-lactamases, enzymes that hydrolyse the *β*-lactam amide, rendering it ineffective in mimicking the cross-linking terminal peptides. The number of identified *β*-lactamases has greatly increased over the years; at the time of writing, the *β*-Lactamase Data-Base [8] shows 8273 identified enzymes.Over the years, extensive use of broad-spectrum *β*-lactams, especially cephalosporins against Enterobacteriaceae since the 1980s, led to selection pressure on bacteria, eventually giving rise to Extended Spectrum *β*-lactamases (ESBLs). Generally, ESBL-producing bacteria are inherently resistant to most penicillins, first-, second-, and third-generation cephalosporins as well as aztreonam. However, most ESBLs typically do not hydrolyse carbapenems, and can be inhibited by some *β*-lactamase inhibitors [4, 9]. Enterobacteriaceae family members have adapted two methods for ESBL evolution: (i) selection of mutants having inherent wide substrate specificity (SHV- and TEM-type- *β*-lactamases) and (ii) acquisition of novel *β*-lactamase genes from the surrounding metagenome [10]. CTX-Ms (cefotaximase) have become the most successful ESBLs of the latter type. They belong to plasmid-encoded Ambler class A active-site serine ESBLs. The most common hosts for this acquired expression of CTX-M type enzyme have been *Escherichia coli* and *Klebsiella pneumoniae*, but it has also been reported in *Salmonella enterica, Shigella spp., Klebsiella oxytoca, Enterobacter spp., Pantoea agglomerans, Citrobacter spp., Serratia marcescens, Proteus mirabilis, Morganella morganii and Providencia spp.* [10]. Out of all the CTX-M sub-lineages, CTX-M-1 and CTX-M-9 clusters are the most widespread. CTX-M-14 (Fig. 1 a) belongs to the CTX-M-9 cluster and is one of the most prevalent members of the group [10]. CTX-M-14 *β*-lactamase hydrolyses most cephalosporins and penicillins as well as carbapenems, although the latter with lower efficiency [11]. Despite their long prevalence, structural information for isoxazolyl antibiotics in complex with Class-A *β*-lactamases is still missing.

**Fig. 1.**
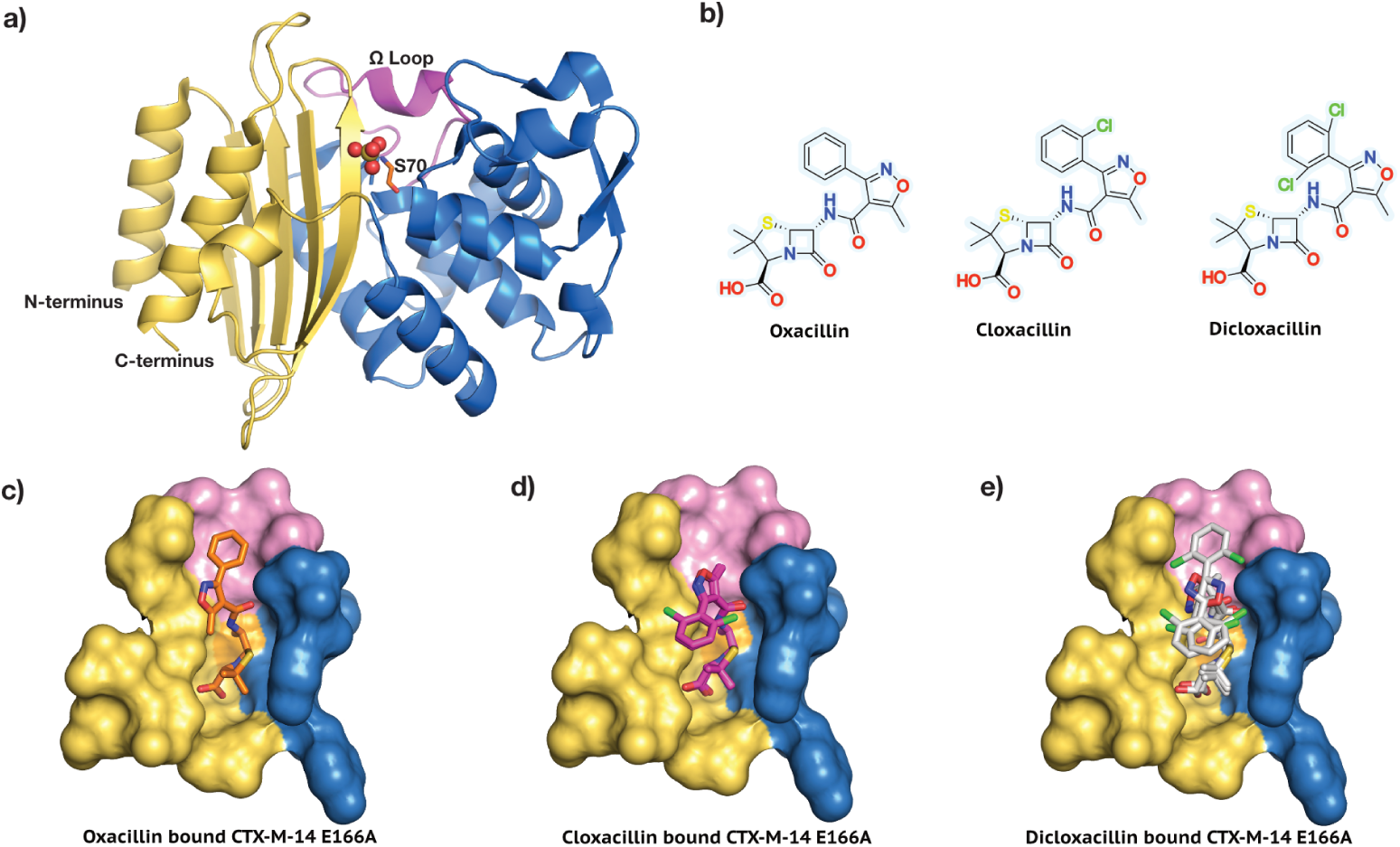
Model system and target ligands. a) CTX-M-14 *β*-lactamase. The catalytically important active site residue Ser 70 and the functionally relevant Ω loop (residues 164-179) are highlighted. b) The isoaxazolyl penicillins, Oxacillin, Cloxacillin and Dicloxacillin are shown in their unhydrolyzed form. c) Oxacillin in the CTX-M-14 E166A active site. d) Cloxacillin in the CTX-M-14 E166A active site. e) Dicloxacillin in the CTX-M-14 E166A active site.

Isoxazolyl penicillins are a class of semi-synthetic antibiotics with effective oral absorption, first produced and tested in the mid-1960s [12, 13]. This class contains four major antibiotics: Oxacillin, Cloxacillin, Dicloxacillin, and Flucloxacillin.

They are synthesised by reacting an isoxazolyl group with the 6-amino-penicillanic acid nucleus derived from penicillin G [14]. These are acid stable and resistant to penicillinase enzymes produced by Gram-positive bacteria. For instance, the therapeutic efficacy of the isoxazolyl penicillins in treating staphylococcal and streptococcal infections was well established in the 1970s [15]. These narrow-spectrum antibiotics, especially Flucloxacillin and Oxacillin, are commonly prescribed against skin and soft tissue infections and osteomyelitis [16]. In an *in-vitro* synergism study against 40 strains of 9 *β*-lactamase producing Gram-negative Enterobacteriaceae members including *E. coli* and *K. pneumoniae*, oxacillin, cloxacillin and dicloxacillin were paired with melzocillin, a semi-synthetic ureido penicillin. The combination was successful for strains of *Morg. morganii, E. coli, Ser. marcescens* but showed no activity against *Klebsiella, Citrobacter or Enterobacter* [17].

To understand the binding mode and the influence of the derivatization in isoxazolyl antibiotics we aimed to gain structural insight into a stable enzyme-substrate complex. As there is currently no structure of a Class-A *β*-lactamase in complex with isoxa- zolyl penicllins available, we utilised the activity-impaired mutant CTX-M-14 E166A. This mutant traps the acyl-enzyme intermediate by covalently binding the isoxazolyl penicillins to the catalytic serine residue (Ser 70). Recent multi-temperature crystallography studies suggests that ambient or physiological temperatures permit resolving a larger conformational ensemble than under cryo-conditions [18, 19, 20, 21]. In other words, data collection closer to physiological temperatures could aid in capturing lower populated conformational states potentially relevant to ligand binding or catalysis. Making use of our recently developed serial crystallography environmental control box [22], we successfully obtained serial room-temperature crystal structures of Oxacillin, Cloxacillin and Dicloxacillin bound to CTX-M-14 E166A (Fig. 1 c-e).

## 2 Results and Discussion

Using room temperature serial synchrotron crystallography (SSX) we have obtained the crystal structures of Class A *β*-lactamase CTX-M-14 mutant E166A, bound to three isoxazolyl penicillins, namely Oxacillin, Cloxacillin and Dicloxacillin (Fig. 2, 3, 4). All crystal structures were obtained at a comparable resolution of 1.8 Å and could be refined to reasonable data-quality parameters (Table 1).

**Fig. 2.**
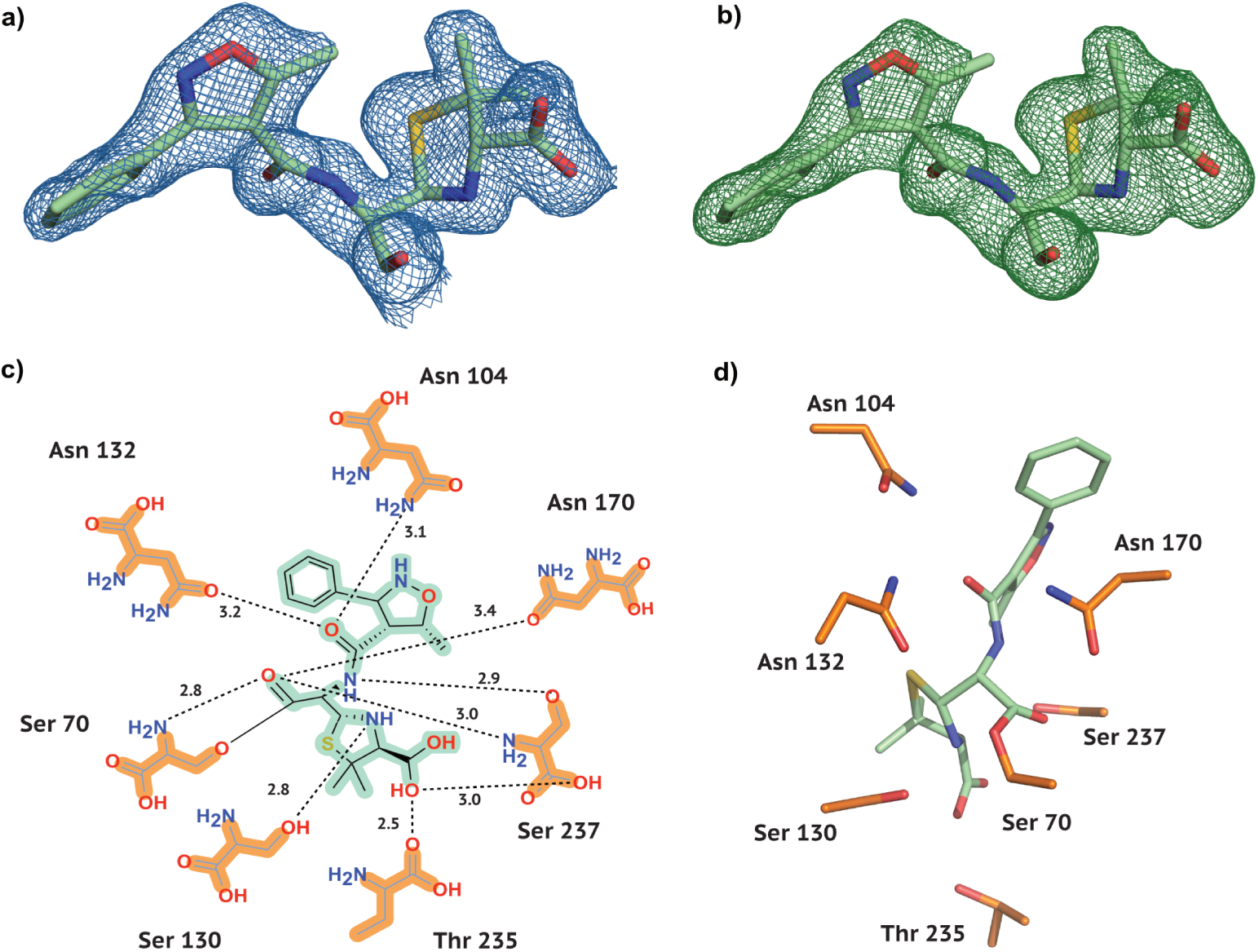
E**l**ectron **density and conformation of Oxacillin at 20 °C** a) 2F*o*-F*c* map shown at an RMSD of 1.0. b) POLDER omit map shown at an RMSD of 3.0. c) 2D projection of Oxacillin and the contact residues at 20 °C showing interatomic distances. d) Oxacillin conformation at 20 °C in the CTX-M-14 active site. Hydrogen bonds are represented as black dotted lines with their distance given in Å.

**Fig. 3.**
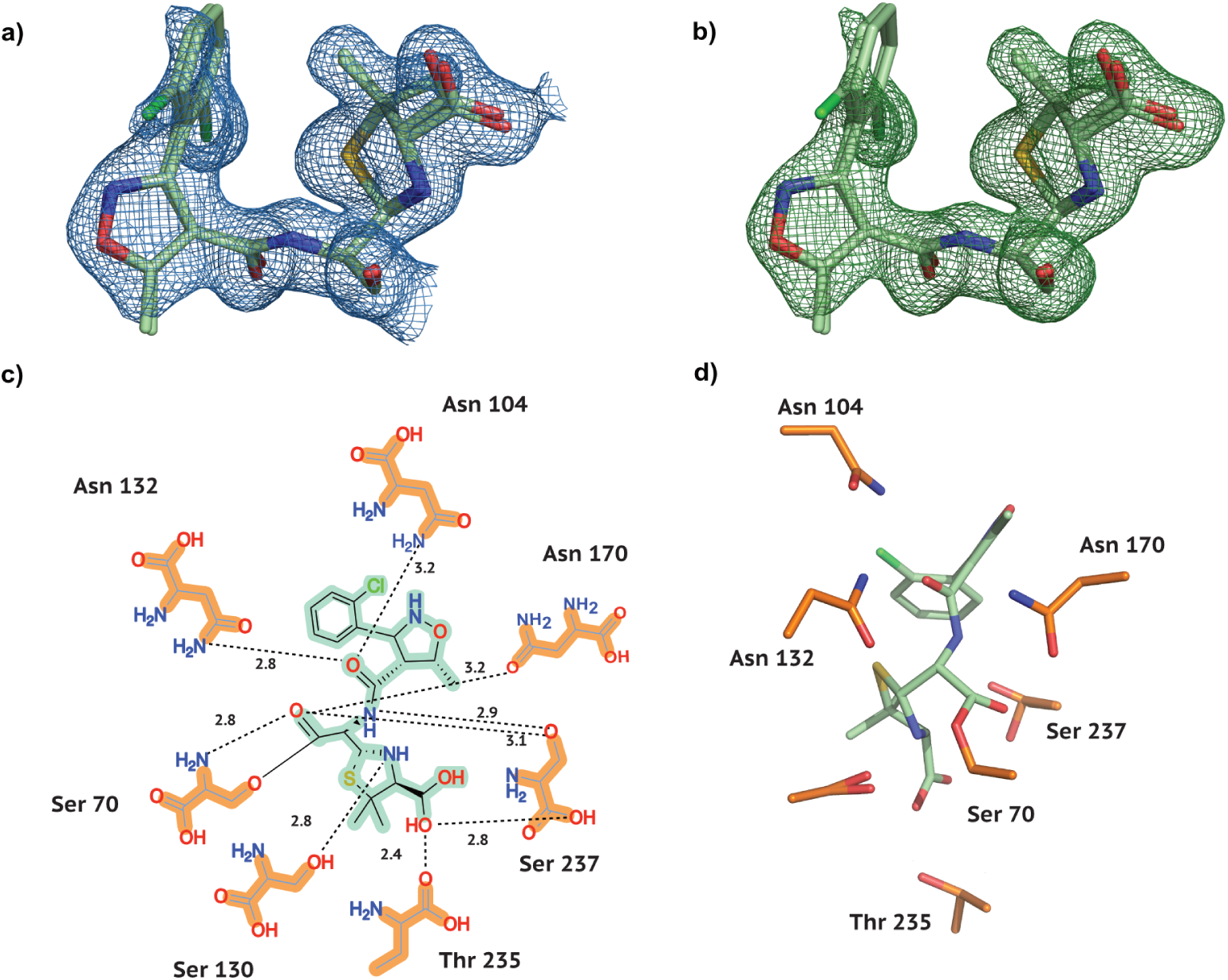
E**l**ectron **density and conformation of Cloxacillin at 20 °C** a) 2F*o*-F*c* map shown at an RMSD of 1.0. b) POLDER omit map shown at an RMSD of 3.0. c) 2D projection of Cloxacillin and the contact residues at 20 °C showing interatomic distances. d) Cloxacillin conformation at 20 °C in the CTX-M-14 active site. Hydrogen bonds are represented as black dotted lines with their distance given in Å.

**Fig. 4.**
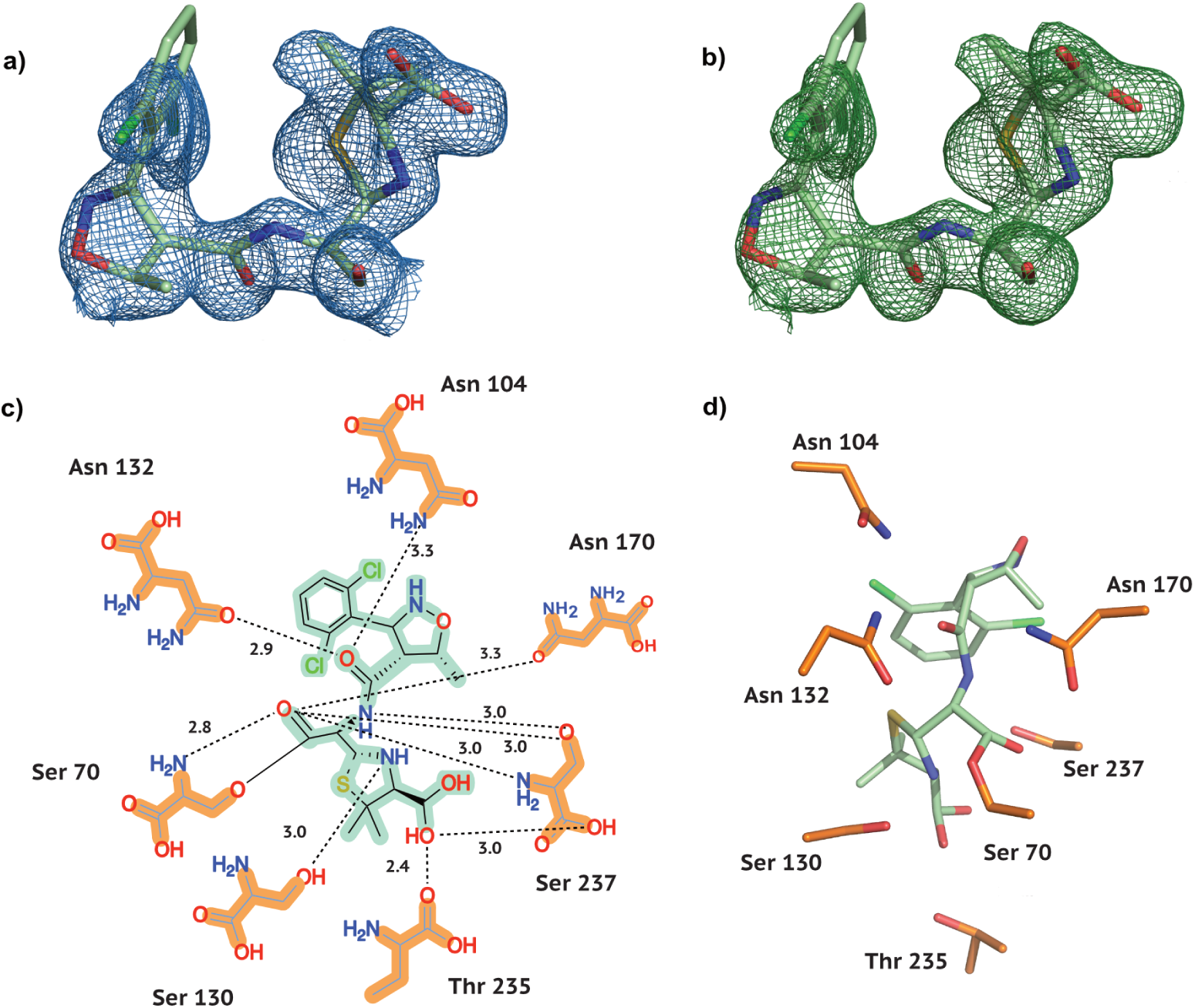
E**l**ectron **density and conformation of Dicloxacillin at 20 °C** a) 2F*o*-F*c* map shown at an RMSD of 1.0. b) POLDER omit map shown at an RMSD of 3.0. c) 2D projection of Dicloxacillin and the contact residues at 20 °C showing interatomic distances. d) Dicloxacillin conformation at 20 °C in the CTX-M-14 active site. Hydrogen bonds are represented as black dotted lines with their distance given in Å.

**Table 1.**
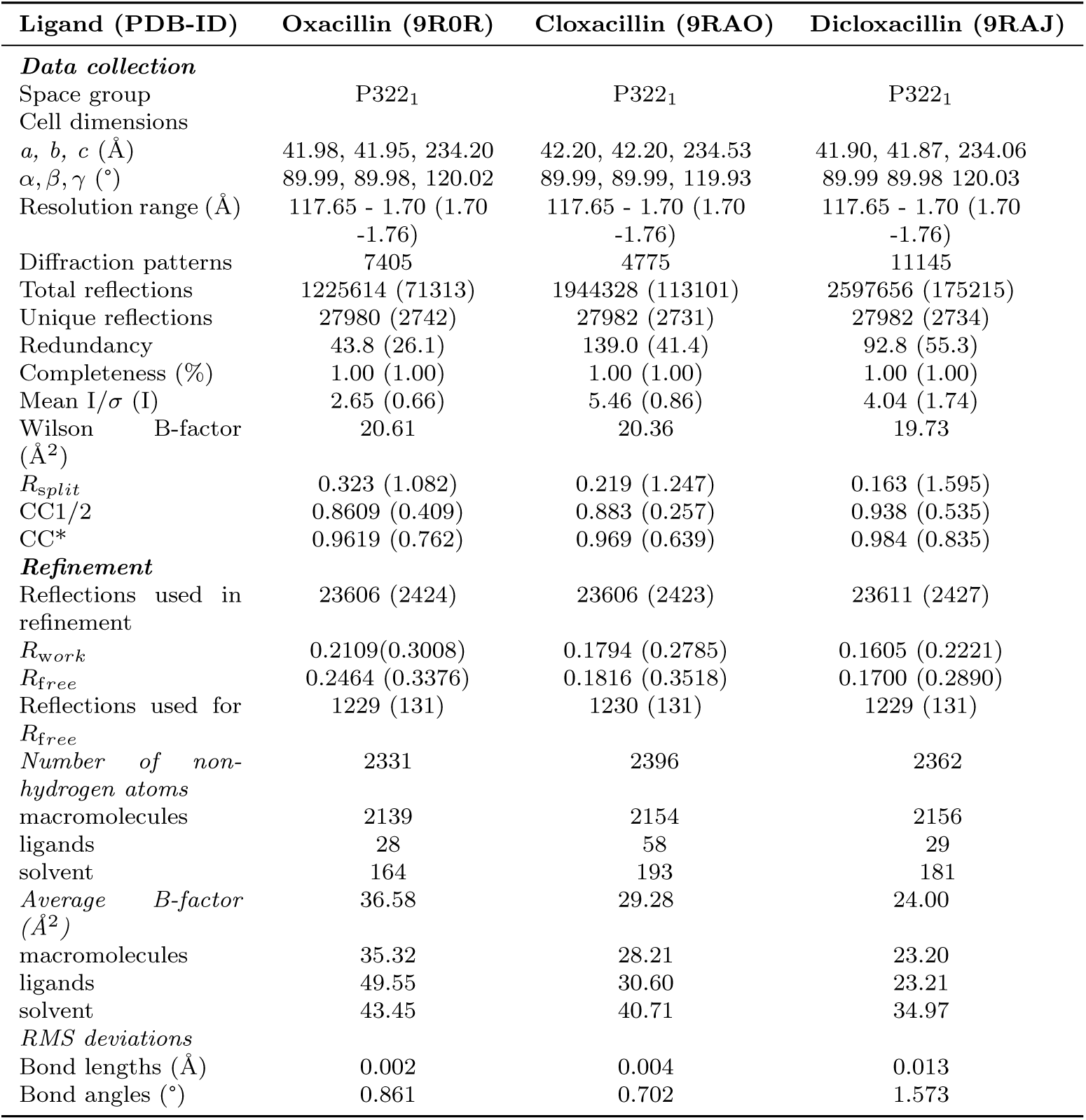
Data collection and refinement statistics of the CTX-M-14 and isoxazolyl complexes at room temperature. Values in the highest resolution shell are shown in parentheses.

As is generally characteristic of penicillin antibiotics, our three substrates con- tain a thiazolide ring attached to a 4-membered *β*-lactam ring (Fig. 1). Furthermore, in isoxazolyl penicillins the amide branch on the *β*-lactam ring opposite to the nitrogen has a variable sidechain, bearing a 3-phenyl-5-methylisoxazole in Oxacillin, a 3-(2-chlorophenyl)-5-methylisoxazole in Cloxacillin, and 3-(2,6-dichlorophenyl)-5- methylisoxazole in Dicloxacillin. After structure solution, successful complex formation with the ligands was clearly evident from the strong difference electron density connected to the catalytic active site residue Ser 70. In addition, typical ligand-residue interactions in the binding pocket of CTX-M-14 were also observed for all three ligands. Commonly, the ligand poses are stabilised by hydrogen bonds between the ligand and the surrounding Ser 70, Asn 104, Ser 130, Asn 132, Asn 170, Thr 235, and Ser 237 residues (Figs. 2,3,4).

In class A *β*-lactamases, the main chain nitrogens of residues Ser 70 and Ser 237 form hydrogen bonds with the carbonyl oxygen of the *β*-lactam; this is thought to stabilize the developing negative charge on the tetrahedral intermediate during the acylation step [23]. This is also observed in all complexes, where 2.8-2.9 Å and 3.0 Å hydrogen bonds are maintained between the *β*-lactam carbonyl and the main chain nitrogens of Ser 70 and Ser 237, respectively.

Previously reported key interactions within the Ω loop (Fig. 1 a) of Class A ESBLs are also observed in all three CTX-M-14 E166A ligand complexes [24, 25], for instance, the H-bonds between Arg 164 and Thr 171, Ala 166 and Asn 136 (contrary to Glu 166 and Asn 170 in the native enzyme). Similarly, salt bridges between Arg 164 and Asp 179, Arg 161 and Asp 163, Asp 176 and Arg 178 are observed. These interactions are vital for maintaining the structural integrity of the Ω loop, which encloses the CTX-M active site. There are no major conformational differences in the Ω loop between the room temperature structures of our three isoxazolyl–CTX-M-14-E166A complexes. Plasmid-born Oxacillin resistance was first mentioned in the late 1960s, and it was hypothesised that this resistance originated from an Oxacillin-hydrolysing *β*-lactamase [26]. Consequently, these enzymes were referred to as Oxacillinases, due to their preferential substrate selection of isoxazolyl-type penicillins. Today, Oxacillinases belong to class D *β*-lactamases and have evolved to become a diverse family with broad substrate profiles including penicillins, extended-spectrum cephalosporins, and even carbapenems [2, 27, 28]. At the time of writing, ten structures of Oxacillin and two of Cloxacillin bound to *β*-lactamases and PBPs are deposited in the PDB. On the other hand, there are only two available structures of Dicloxacillin in the PDB, and neither is bound to a *β*-lactamase or PBP. Notably, there are no PDB entries of the isoxazolyl penicillins in complex with a class A *β*-lactamase. Furthermore, all deposited structures are collected at cryo-temperatures (Table 2).

**Table 2.** List of PDB entries with Oxacillin, Cloxacillin, or Dicloxacillin, respectively.

| Ligand | Class B | Class C | Class D | PBPs | Non-β-lactamases |
| --- | --- | --- | --- | --- | --- |
| Oxacillin | 4EYB | 4JXG<sup>†</sup> | 4F94, 7O5Q<sup>†</sup>,<br>7PSE, 4MLL | 8VUB, 5OJ1,<br>5OIZ, 5M19 | -<br>- |
| Cloxacillin | - | 1FCM | - | 3MZD | - |
| Dicloxacillin | - | - | - | - | 4R23, 6ZOA |

<sup>†</sup> unpublished structures

### 2.1 Comparison of the Oxacillin complex to the OXA-I Class D *β*-lactamase

To compare our room-temperature Oxacillin-bound CTX-M-14-E166A complex (Fig. 2) to previously solved Oxacillin complexes, we selected the class D *β*-lactamase structure OXA-1 (Table 2, PDB entry 4MLL, Fig. 5) [28]. OXA-1 is an important subtype of Ambler Class D enzymes, which effectively hydrolyse Oxacillin. Its 3D structure was previously solved at a resolution of 1.4 Å, and it shares 16.5% sequence identity with CTX-M-14-E166A, while the other published class D enzymes in complex with Oxacillin, 7PSE [29] and 4F94 [28] share only 6.4% and 13.5% sequence identity, respectively. Since the Class B New Delhi metallo-*β*-lactamase (PDB entry 4EYB) displays a different binding mode and lower homology to CTX-M-14 (13.2% sequence identity), this structure as well as the unpublished Class C and Class D entries were not considered for a detailed comparison [30].

**Fig. 5.**
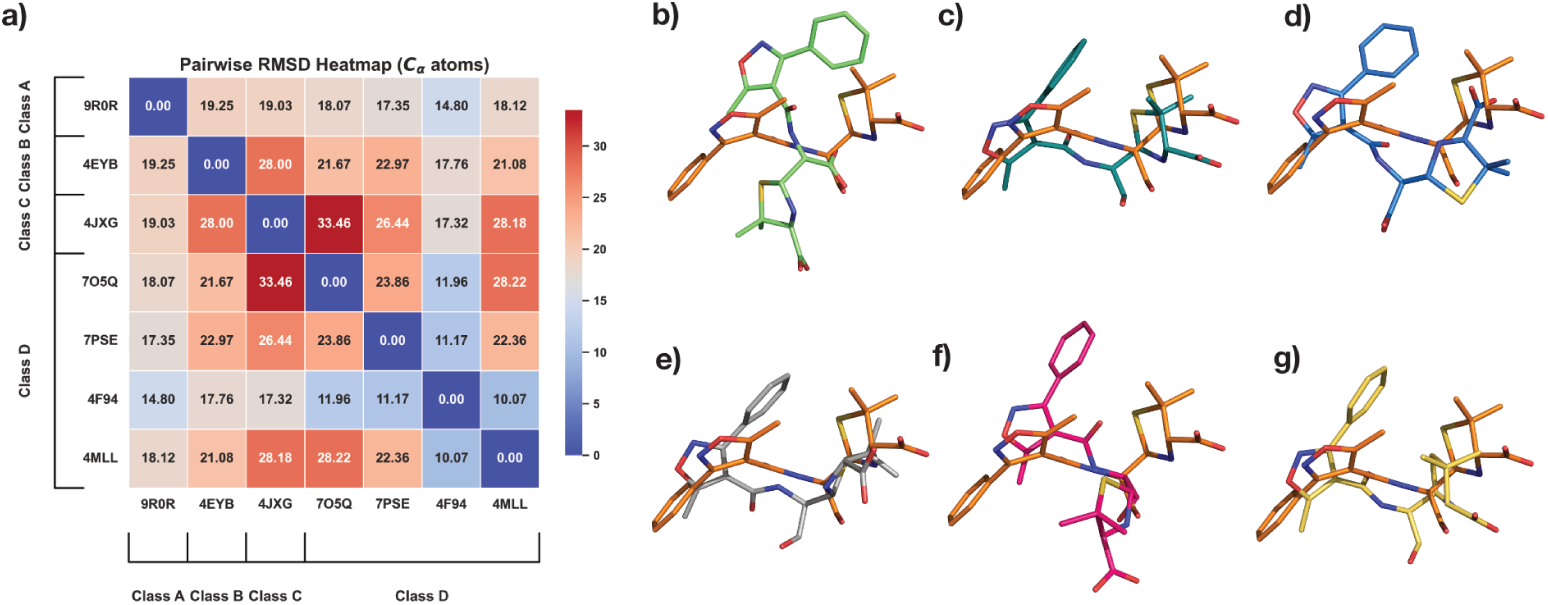
P**r**otein **alignment and Oxacillin superposition** a) Pairwise C*α* heatmap for all Oxacillin bound *β*-lactamase complexes available in the PDB. Figures b-g show CTX-M-14 E166A bound Oxacillin, PDB entry 9R0R,(orange) superposed to Oxacillin from PDB entry b) 4EYB with an RMSD of 5.45 Å, c) 4JXG with an RMSD of 4.51 Å, d) 7O5Q with an RMSD of 5.02 Å, e) 7PSE with an RMSD of 4.51 Å, f) 4F94 with an RMSD of 5.23 Å, g) 4MLL with an RMSD of 4.46 Å.

OXA-1 (4MLL) has four monomers out of which 3 show bound Oxacillin at the active site. Monomer C was chosen for comparison, having lowest B factors as well as least amount of difference density features in the accompanying electron density map. In OXA-I, the carboxyl oxygen of the *β*-lactam ring forms H-bonds with the main-chain nitrogens of Ala 215 and active site Ser 67, which correspond to Ser 237 and active Ser 70 in CTX-M-14. Along the same lines, the amide nitrogen of the variable side-chain moiety forms an H-bond with the backbone carbonyl of Ala 215 in OXA-1, while in CTX-M-14 it is connected to Ser 237. Usually, the amide oxygen interacts with a polar residue (Asn 104 and Asn 132 in CTX-M-14) in *β*-lactamases. However, in OXA-1, a hydrogen bond with a water molecule is observed, due to the hydrophobic Val 117 in place of a polar residue. The carboxylate of the thiazolide ring forms hydrogen bonds with Ser 258, Thr 213 and waters in OXA-1, while the polar contacts to water molecules vary between the four OXA-I monomers in the asymmetric unit. In CTX-M-14 on the other hand, Thr 235 and Ser 237 interact with the thiazolide ring. It has been reported that class D enzymes possess a more hydrophobic active site than class A enzymes [31], which complements the largely hydrophobic moiety of Oxacillin [28]. Nevertheless, we have observed a stable Class A *β*-lactamase–Oxacillin complex with unambiguous electron density, further underlining the broad substrate spectrum of class A ESBLs. To illustrate localised difference in the binding mode of Oxacillin with different *β*-lactamases, we superposed our CTX-M-14 E166A bound Oxacillin with all other *β*-lactamase-bound Oxacillin molecules in the PDB (Fig. 5 b-g). Comparing the C*_α_* RMSD (Fig. 5 a) with the superposed ligands (Fig. 5 b-g) side by side does not reveal a direct correlation between enzyme similarity and resemblance in lig- and binding mode. Interestingly, in contrast to all previously studied complexes, our room-temperature structure reveals a markedly different binding pose for Oxacillin. While the 3-phenyl-5-methylisoxazole moiety is oriented toward Pro 167 of CTX-M-14 E166A (Fig. 5 b–g), this group is rotated by more than 150° in all other oxacillin- bound structures. Whether this difference is rooted in the structural differences of Class A *β*-lactamases or can be attributed to the higher data-collection temperature will be explored in future studies.

### 2.2 Comparison to other Cloxacillin complexes

To compare our Cloxacillin complex to previously solved structures, we selected the complex with Class C *β*-lactamase AmpC [32] (PDB entry 1FCM, 2.46 Å resolution, 14.5% sequence identity) from *E. coli*, and the *E. coli* Penicillin-Binding Protein 5 [33] (PDB entry 3MZD, 1.90 Å resolution, 12.2% sequence identity). Cloxacillin-PBP-5 forms multiple H-bonds between the carboxyl of the thiazolide ring and residues Arg 248 and Thr 214. Additionally, the main chain nitrogens of the active Ser 44 and Gly 215 form H-bonds with the carboxyl oxygen of the *β*-lactam ring. The amide branch of the *β*-lactam ring, to which the variable moiety is attached, forms multiple H-bonds with Ser 87 and Asn 112. Along similar lines, in the Cloxacillin-AmpC structure, the thiazolide carboxyl is stabilised by multiple polar interactions with Thr 313, Asn 343, and multiple water molecules. Active site Ser 61 and Ala 315 form hydrogen bonds with the *β*-lactam carboxyl. Finally, the amide oxygen interacts with Asn 149 and the thiazolide ring nitrogen with a water molecule. In our Cloxacillin complex (Fig. 3 c,d), a few parallels can be drawn; for example, the thiazolide carboxyl forms H-bonds with side chain oxygens of Thr 235 and Ser 237. The amide branch of the *β*-lactam ring forms multiple H-bonds with Asn 104, Asn 132, and Ser 237. Additionally, the ligand is also stabilised by H-bonds between Asn 170 and the carboxyl oxygen of the *β*-lactam and that of Ser 130 with the nitrogen of the thiazolide ring. Within PBP-5, most of the electron density corresponding to the thiazolidine ring and carboxylate of Cloxacillin is clearly resolved, except for one of the two methyl groups at position 2 of the ring [33]. The presence of weak electron density around the variable moiety, especially for the 2-chlorophenyl ring, has previously been reported [33, 32]. In contrast, we observe clear and strong density peaks surrounding the whole ligand except the 2-chlorophenyl ring (Fig. 3 a,b). This might be a result of rotation on the 2-chlorophenyl ring about the C-C bond connecting it to the oxazole. To justify strong positive density peaks around the oxazole, an additional alternate conformation of the variable moiety was modelled. As a result of occupancy refinement, the major conformer has 75% and the minor conformer has 25% occupancy. Only the major conformer is referred to in the analysis and Fig. 3,6. Superposing our Cloxacillin complex with *E. coli* PBP-5 (3MZD) [33] and the Class C *β*-lactamase AmpC [32] (1FCM) (Fig. 7) highlights the marked differences in the ligand conformations with respect to the enzymes.

**Fig. 6.**
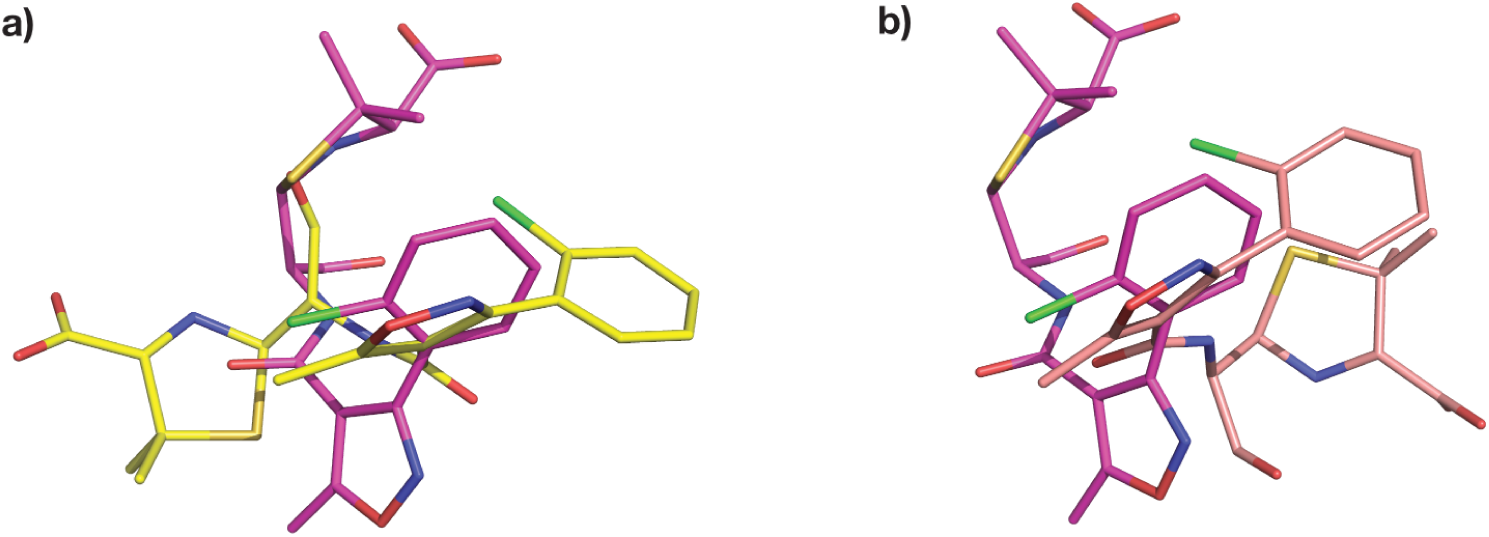
C**l**oxacillin **superposition** a) CTX-M-14 E166A bound Cloxacillin,PDB entry 9RAO, (magenta) superposed to Cloxacillin from PDB entry 1FCM (yellow) with an RMSD of 3.98 b) CTX- M-14 E166A bound Cloxacillin,PDB entry 9RAO, (magenta) superposed to Cloxacillin from PDB entry 3MZD (salmon) with an RMSD of 4.68.

**Fig. 7.**
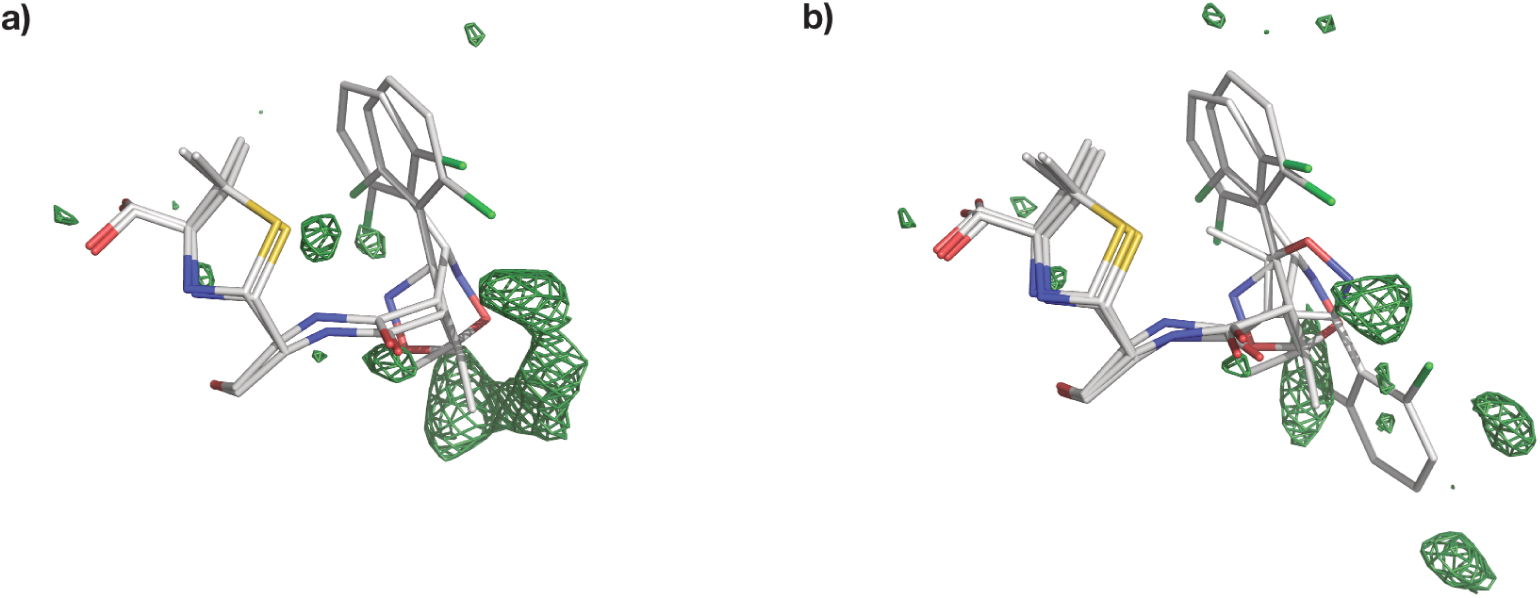
M**u**lti**-conformer structure of Dicloxacillin bound to CTX-M-14 E166A** a) One alternate conformation along with F*o*-F*c* map around 3Åof the ligand at an RMSD of +3.0 b) A third conformation reduces some of the persistent difference density from a); F*o*-F*c* map around 3Åof the ligand at an RMSD of +3.0

### 2.3 Dicloxacillin binding mode

To the best of our knowledge, our CTX-M-14 E166A Dicloxacillin complex is the first observed 3D structure of the antibiotic with a *β*-lactamase enzyme. Dicloxacillin displays an unambiguous difference electron density (Fig. 4 a,b) and clearly binds to CTX-M-14 E166A. A careful observation of the difference map indicates the pos- sibility of at least two more alternate conformations as a result of rotation on the 2-chlorophenyl ring about the C-C bond connecting it to the oxazole. Thus, two orientations are similar to Cloxacillin with of 40% and 21% occupancies. Adding a third ring orientation (Fig.7 a,b) reduces some of the strong positive difference electron density around the ligand. However, this conformer, with an occupancy of 39% differs from Cloxacillin and resembles Oxacillin instead (Fig. 7 b,c). Such a multi-conformer Dicloxacillin model is strongly supported by the F*_o_*-F*_c_* map as well as the POLDER omit map. Nevertheless, only a single ligand orientation (40% occupancy) is represented in Fig. 4. This Dicloxacillin conformer also interacts with the binding pocket via residues mentioned above (Fig. 4 c,d). The alternate, almost equally occupied (39%) Oxacillin resembling conformer, leads to additional ligand-residue interactions, notably, the 5-chlorine interacts with the backbone nitrogen of Asp 239, backbone oxygen of Asn 170 and the sidechain nitrogen of Asn 104.

### 2.4 Isoxazolyl Penicillin binding to Class A *β*-lactamases

We have obtained reliable structures of Oxacillin, Cloxacillin and Dicloxacillin bound to CTX-M-14 E166A mutant at room temperature, displaying strong electron density around the covalently bound isoxazolyl penicillins. Interestingly, Oxacillin does not adopt an alternative conformation at 20 °C. The electron density around Oxacillin clearly indicates the presence of only one orientation and leaves no room even for a very low occupancy alternate conformer. On the other hand, Cloxacillin and Dicloxacillin clearly adapt multiple conformations as mentioned in previous sections. The 2-chlorophenyl ring of Cloxacillin assumes one additional orientation, whereas the 2,5-dichlorophenyl ring of Dicloxacillin shows at least two alternate orientations. There are still observable minor blobs of positive and negative electron density surrounding Dicloxacillin, signatures of its partially dynamic binding mode. To avoid overfitting, we have left these minor populated states unmodelled. Unlike Oxacillin, the presence of chlorobenzenes in the latter two ligands may point towards their dynamicity at room temperature. This may also exemplify gradual and subtle changes in ligand binding between the lead compound (Oxacillin) and its derivatives (Cloxacillin, Dicloxacillin). Quite interestingly, as Fig. 8 indicates, Dicloxacillin seems to adapt conformations resembling both Cloxacillin and Oxacillin. As described in the previous section, 5-chlorine of Dicloxacillin additionally interacts with neighbouring residues, leading to one additional orientation compared to Cloxacillin. How much of this conformational diversity between the present isoxazolyl penicillins pertains only to the phenyl chlorination and to the temperature will be systematically explored in further studies.

**Fig. 8.**
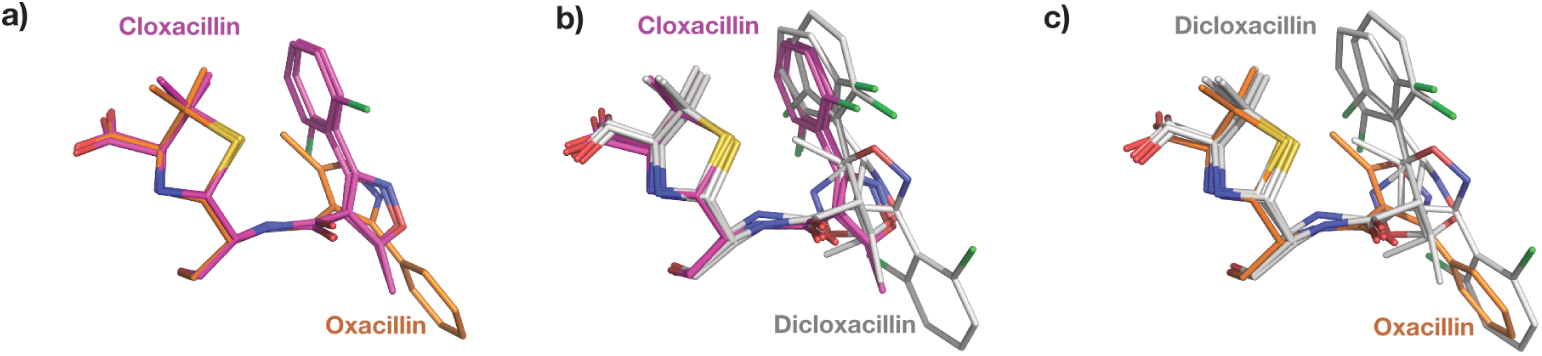
S**u**perposed **Isoxazolyls bound to CTX-M-14 E166A** a) Oxacillin (Orange) superposed with Cloxacillin (Magenta) b) Cloxacillin (Magenta) superposed with Dicloxacillin (Grey) c) Dicloxacillin superposed with Oxacillin

### 2.5 Outlook

With the advent of methodologies like serial synchrotron crystallography (SSX), data collection close to or at physiological temperatures has become more accessible to a larger user community, as it enables effectively mitigating radiation damage [34, 35, 36]. Consequently, protein-ligand interaction studies can become more insightful. Room temperature data collection removes vitrification artefacts and ice formation, enabling observations of conformational heterogeneity. Despite a somewhat lower throughput than other sample delivery methods, fixed-target SSX is very material efficient and reduces radiation damage effects by distributing the dose over thousands of crystals [34, 35, 36, 37, 38]. Additionally, our environment control system provides constant high relative humidity and prevents non-isomorphism caused by dehydration [22]. Collectively, these aspects mitigate cryo-crystallography artefacts, and allow to study enzyme-ligand interactions, e.g. for structure-based drug design. Nevertheless, SSX’s potential in the realm of drug discovery remains an avenue that has not yet been extensively explored. Recent studies have utilized serial crystallography for fragment screening and ligand binding campaigns [39, 40, 41]. The authors have successfully reported additional binding sites as well as protein allosteric conformational responses, emphasizing the benefits of room-temperature crystallography for studies aiming to support structure-based drug design. On similar grounds, have aimed to dive into the physiologically relevant study of existing substrate candidates. As quoted previously, Oxacillin, Cloxacillin and Dicloxacillin have been a part of a synergy study [17] for Gram-negative bacterial strains. Moreover, Dicloxacillin was almost consistently associated with the lowest Minimum Inhibitory Concentration (MICs) attainable for Mezlocillin in combination. This knowledge, combined with the structural insights presented here, could open avenues for new inhibitor and antibiotic designing as well as rigorous synergy studies involving the isoxazolyl penicillins, specifically for the most ubiquitous *β*-lactamase CTX-M-14, as well as other ESBLs.

## 3 Materials and Methods

### 3.1 Protein purification and sample preparation

The CTX-M-14 E166A gene was synthesized and cloned in a pET-24a(+) vector (Bio- Cat GmbH, Heidelberg, Germany) with a Kanamycin selection marker. Expression and purification were performed as described previously [42]. For crystallisation, the CTX-M-14 E166A solution (22 mg/ml) was mixed with 45% (v/v) crystallising agent (40% (w/v) PEG 8000, 200 mM LiSO_4_, 100 mM sodium acetate, pH 4.5) and with 5% (v/v) undiluted seed stock solution to induce micro-crystallization. This resulted in crystals with a homogeneous size distribution of ca. 11-15 µm overnight. For soaking experiments, Oxacillin, Cloxacillin and Dicloxacillin were dissolved in the stabilization buffer (28% (w/v) PEG 8000, 140 mM LiSO_4_, 70 mM sodium acetate, 6 mM MES pH 4.5) to reach a solubility of about 500 mM each. The ligands did not dissolve completely; hence, the supernatant was mixed with the crystal slurry in a 3:1 ratio. Our efforts to soak Flucloxacillin into CTX-M-14 E166A crystals, unfortunately, remained unsuccessful.

### 3.2 Ambient temperature data collection

Serial diffraction data were collected at the EMBL endstation P14.2 (T-REXX) at the PETRA-III synchrotron (DESY, Hamburg, Germany) with an X-ray beam of 10 *×* 7 µm (H*×*V) on an Eiger 4M detector (Dectris, Baden-Daettwil, Switzerland). Before sample loading, the HARE-chip was glow-discharged to increase surface hydrophilicity and thus, improve sample-loading homogeneity [43]. Roughly 120 *µ*l ligand-soaked microcrystal slurry (90 *µ*l ligand solution mixed with 30 *µ*l crystal slurry) was then pipetted onto the fixed target HARE-chip containing 20,736 wells, in a humidified (98%) environment. Data collection was conducted as previously described [44] within the environmental control box [22] mounted on the T-REXX endstation. Briefly: After mounting, the chips were raster-scanned through the X-ray beam using a 3-axis piezo translation stage setup (SmarAct, Oldenburg, Germany) [43]. Data collection was carried out at 20 °C with the relative humidity controlled at 98%. For reasonable data statistics approximately 5000 to 11000 still diffraction images were recorded per structure as previously determined as a broad guideline [34, 45].

### 3.3 Data-processing and structure refinement

Diffraction data were processed using the CrystFEL v0.09.1 package [46]. Structures were solved by molecular replacement using PHASER with our previously determined CTX-M-14 structures as a search model with one molecule in the asymmetric unit (PDB-ID: 6GTH) [47]. Structures were refined with iterative cycles of *phenix.refine* and manual model building in COOT-v0.9 [48, 49, 50]. Figures were generated with PyMOL [51] and ChemDraw (Revvity Signals Software, Inc., Waltham, USA). Data collection and refinement statistics are summarized in Table 1.

### 3.4 Data analysis

To illustrate localised differences in ligand bidning, the structures were superimposed with the PyMOL function pair_fit, which allowed comparison of the ligand and the binding pocket without outlier rejection. Fig. 5a visualises C*_α_* RMSD of all Oxacillin- bound *β*-lactamases from Table 2 as a categorical heatmap. For each pair, the two structures were aligned with PyMOL function align to minimise the resulting RMSD value. The heatmap was generated with the resulting RMSD values using python module matplotlib.

## Acknowledgments

All ambient-temperature SSX data were collected at beamline P14.2 (T-REXX) operated by EMBL Hamburg at the PETRA-III storage ring (DESY, Hamburg, Germany). We would like to thank our colleagues A.R. Pearson and P. Mehrabi for their continuous support and helpful discussions. We are grateful to Hamid Nasiri for the kind provision of Cloxacillin and Dicloxacillin.

## Author Contributions

E.C.S. designed the experiments; G.G. and E.C.S. performed the data collection with support from D.v.S.; K.B. and A.P. prepared the protein; A.P. prepared the protein crystals; G.G., D.v.S. and E.C.S. processed and analysed the diffraction data. G.G. refined the structures. G.G. and E.C.S. wrote the manuscript; All authors discussed and corrected the manuscript.

## Declarations

Competing Financial Interests Statement: The authors declare no competing financial interests.

